# Ancient coding sequences underpin the spatial patterning of gene expression in C_4_ leaves

**DOI:** 10.1101/085795

**Authors:** Ivan Reyna-Llorens, Steven J. Burgess, Ben P. Williams, Susan Stanley, Chris Boursnell, Julian M. Hibberd

**Affiliations:** Department of Plant Sciences, Downing Street, University of Cambridge Cambridge CB2 3EA, UK.

## Abstract

Photosynthesis is compromised in most plants because an enzymatic side-reaction fixes O_2_ instead of CO_2_. The energetic cost of oxygenation led to the evolution of C_4_ photosynthesis. In almost all C_4_ leaves compartmentation of photosynthesis between cells reduces oxygenation and so increases photosynthetic efficiency. Here we report that spatial expression of most C_4_ genes is controlled by intragenic *cis*-elements rather than promoter sequence. Two DNA motifs that cooperatively specify the patterning of genes required for C_4_ photosynthesis are identified. They are conserved in plants and algae that use the ancestral C_3_ pathway. As these motifs are located in exons they represent duons determining both gene expression and amino acid sequence. Our findings provide functional evidence for the importance of transcription factors recognising coding sequence as previously defined by genome-wide binding studies. Furthermore, they indicate that C_4_ evolution is based on ancient DNA motifs found in exonic sequence.

## Introduction

Photosynthesis allows atmospheric CO_2_ to be fixed into organic molecules and therefore forms the basis of life on the planet. When plants moved onto land they inherited the photosynthetic system first developed by bacteria, in which the enzyme Ribulose Bisphosphate Carboxylase Oxygenase (RuBisCO) generates the three-carbon compound phosphoglyceric acid (PGA) (Anbar et al., 2007). As PGA contains three carbon atoms, this form of photosynthesis is known as the C_3_ pathway. However, a side-reaction of RuBisCO fixes O_2_ rather than CO_2_, and this generates the toxic compound phosphoglycolate. Although plants use the photorespiratory pathway to remove phosphoglycolate, it is energetically expensive and some carbon is lost in the process (Bauwe et al., 2010). Around 30 million years ago, some plants evolved a photosynthetic system in which CO_2_ is concentrated around RuBisCO such that oxygenation is minimised, and so photosynthetic efficiency increases by around 50% (Hatch and Slack 1966; Sage et al., 1999). These species now represent the most productive vegetation on the planet (Sage et al., 2004; Ray et al., 2012), and because they initially generate a C_4_ acid in the photosynthetic process, are known as C_4_ plants.

The mechanism by which CO_2_ supply to RuBisCO is increased in C_4_ species depends on the spatial separation of photosynthetic reactions. Initial production of C_4_ acids takes place in one compartment, and then their re-release to concentrate CO_2_ occurs in another. Although in some species this can take place within a single cell (Edwards et al., 2004), in the majority of C_4_ plants, evolution has co-opted the existing compartmentation afforded by multi-cellularity to separate these carboxylation and decarboxylation reactions (Hatch and Slack 1966). The separation of photosynthesis between cells requires the co-ordinated regulation of numerous enzymes and transporters. For example, in mesophyll (M) cells, enzymes such as carbonic anhydrase, phospho*eno*/pyruvate carboxylase and malate dehydrogenase allow the production of C_4_ acids from HCO_3_^−^, whereas in the bundle sheath (BS), high activities of C_4_ acid decarboxylases and RuBisCO allow efficient entry of carbon into the Calvin-Benson cycle (Furbank, 2011). The importance of each enzyme accumulating in the correct cell-type is considered critical for the efficiency of the pathway, and so the spatial patterning of photosynthesis gene expression in C_4_ plants has received significant attention. As with most other eukaryotic systems, although post-transcriptional and translational regulation are acknowledged (Hibberd and Covshoff 2010; Kajala et al., 2011; Williams et al., 2016; Fankhauser and Aubry, 2016), most analysis has focussed on the importance of promoters in regulating gene expression in M or BS cells (Sheen, 1991; Viret et al., 1994; Taniguchi et al., 2000; Nomura et al., 2000; Kaush et al., 2001; Gowik et al., 2004; Akyildiz et al., 2007). Previously, we found that expression of multiple *NAD-dependent MALIC ENZYME (NAD-ME)* genes in the BS of C_4_ *Gynandropsis gynandra* was dependent on sequences not found in promoter elements upstream of these genes, but rather in exonic sequence within the gene (Brown et al 2011). Whilst the exact *cis*-elements were not defined, orthologous genes from the C_4_ model *Arabidopsis thaliana* also contained the regulatory DNA necessary for preferential expression in BS cells of the C_4_ leaf. Overall, these data implied that evolution has repeatedly made use of pre-existing regulatory DNA found within genic sequence to pattern gene expression in C_4_ leaves. However, the specific sequences responsible for preferential expression in the BS were not defined, and so it was not clear if exactly the same *cis*-elements were used by each gene. Furthermore, without a definition of the motifs specifying expression in the BS contained within these *NAD-ME* genes, it was not possible to identify if they control expression of additional genes, nor to understand if these same elements are used in other species. To address this, we first identified the sequence motifs responsible for patterning these *NAD-ME* genes, and in doing so, show that they act as duons impacting both on patterns of gene expression as well as amino acid sequence of the encoded protein. We used these sequence motifs to predict and validate other genes that are also controlled by these elements, and also to investigate the likely origin of such regulation within land plants. We report both widespread use of, and ancient origins for, two *cis*-elements that act within coding sequence co-operatively to generate BS expression in C_4_ leaves. We place these findings in the context of genome-wide studies reporting the widespread binding of transcription factors to genic sequence rather than promoter elements, as well as the polyphyletic evolution of the complex C_4_ pathway.

## Results

### Two motifs within coding sequence act co-operatively to generate BS specificity

To better understand the mechanism responsible for generating preferential gene expression in BS cells analysis was focussed on a region of 240 nucleotides within the coding region of *GgNAD-ME1.* Although previous work showed this fragment confers BS accumulation of GUS in *G. gynandra* (Brown et al., 2011) it was unclear if this was due to recognition of the DNA or RNA sequence. To test if regulation was lost when a complementary mRNA was produced, an antisense construct for this 240 nucleotide sequence (Figure 1A) was placed under control of the constitutive CaMV35S promoter, and fused to *uidA* encoding the β-glucoronidase (GUS) reporter. Whereas the CaMV35S promoter alone lead to similar accumulation of GUS in M and BS cells (Figure 1B), microprojectile bombardment of the antisense construct maintained preferential accumulation in the BS (Figure 1B) indicating that DNA sequence is recognised by *trans*-acting factors and therefore that preferential accumulation in BS cells is regulated during transcription.

**Figure 1:**
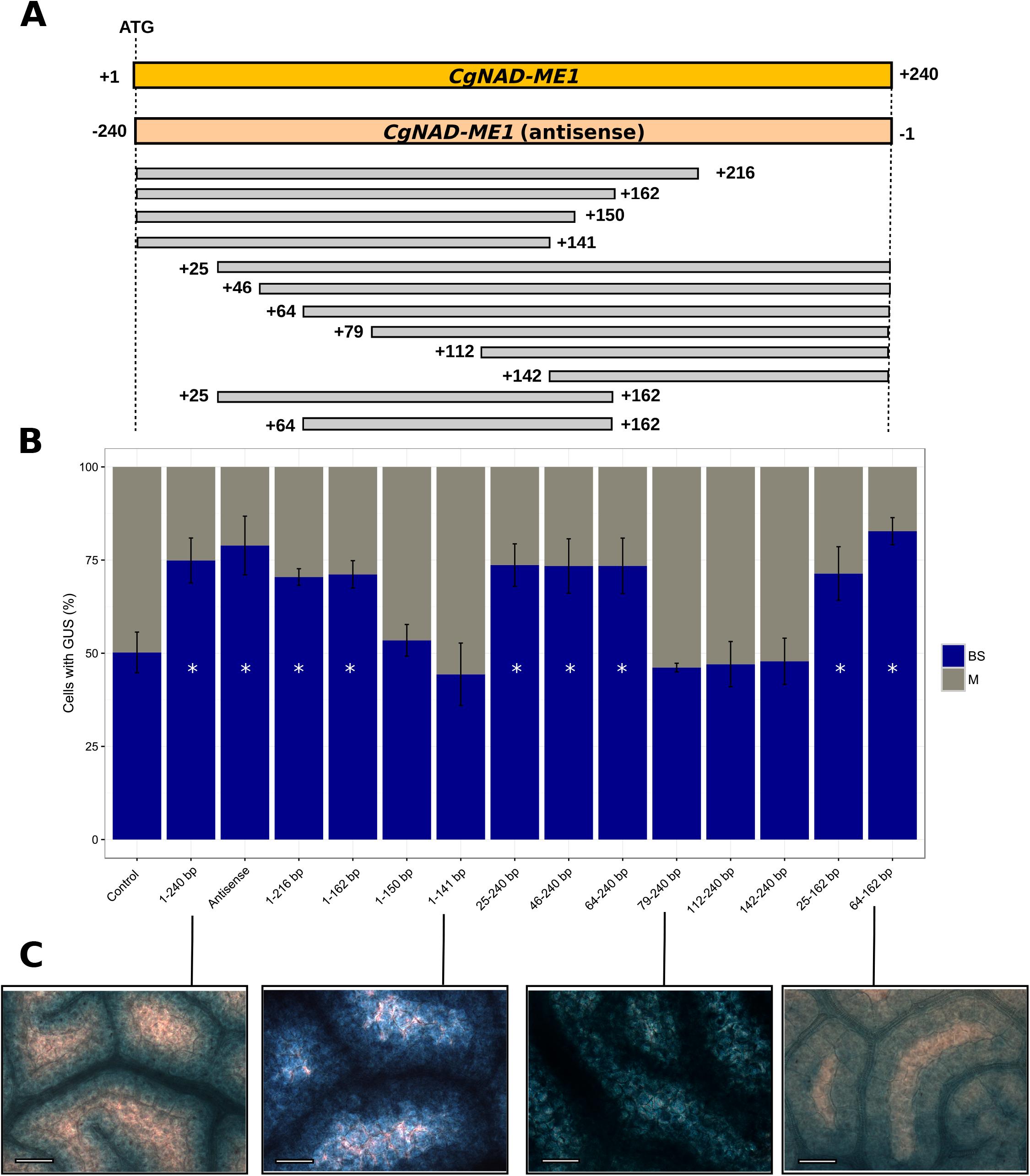
**Two regions within the coding sequence of *GgNAD-ME1* are necessary for preferential gene expression in the bundle sheath.** An antisense construct, as well as a deletion series from the 5’ and 3’ ends of *GgNAD-ME1*_*(1-240 bp)*_ coding sequence were translationally fused to the *uidA* reporter under the control of the CaMV 35S promoter **(A).** Percentage of cells containing GUS after microprojectile bombardment of *G. gynandra* leaves. Bars represent the percentage of stained cells in BS (blue) and M (grey) cells, error bars denote the standard error. * represents statistically significant differences with P-values <0.05 and CI = 95% determined by a one-tailed t test **(B).** GUS in *G. gynandra* transformants containing uidA fused to 1-240, 1-141, 79-240 and 64-162 bp from the translation starting site of *GgNAD-ME1* **(C).** Scale bars, 100 µm.

To identify the specific nucleotides responsible for BS expression within this 240bp fragment a deletion series was generated by removing fragments from either the 5’ or 3’ end of *GgNAD-ME1* (Figure 1A). Deletion of 24, and 78 nucleotides from the 3’ end did not affect preferential accumulation in BS cells (Figure 1A-B), but removal of 90 nucleotides resulted in loss of cell specificity (Figure 1A-C). Similarly, deletion of the first 63 nucleotides from the 5’ end did not abolish preferential accumulation of GUS in BS cells, but removing 78 nucleotides did (Figure 1A-C). A fragment incorporating bases 64 to 162 was sufficient to retain cell preferential accumulation in the BS both after microprojectile bombardment (Figure 1B), and after production of stable transformants (Figure 1C, S2). We conclude that one region composed of the nucleotides TTGGGTGAA (64 to 79 downstream of the translational start codon) and a second region made up of nucleotides GATCCTTG (141 to 162 nucleotides downstream of the translational start codon) are necessary for preferential accumulation of *GgNAD-ME1* in BS cells of C_4_ *G. gynandropsis.* These two regions will hereafter be referred to as Bundle Sheath Motif 1a (BSM1a) and Bundle Sheath Motif Ib (BSM1b), and they are separated by 75 nucleotides.

To test if the sequence separating BSM1a and BSM1b is required for preferential expression in BS cells, it was replaced with exogenous sequence lacking homology to the native region of *GgNADME1* (Figure 2A). The fragment that contained BSM1a and BSM1b separated by this exogenous sequence led to preferential accumulation of GUS in BS cells (Figure 2A, 2B). Although the exact sequence separating BSM1a and BSM1b does not impact on their function, the distance separating them could play an important role. The length of the spacer was therefore modified, and this indicated that BSM1a and BSM1b do not generate preferential accumulation in the BS cells when fused together directly, or when separated by 999 base pairs (Figure S3). However, when the intervening sequence was between 21 and 550 base pairs preferential accumulation in BS cells occurred (Figure S3). Site-directed mutagenesis of each motif showed that the first two nucleotides of BSM1a had no impact on preferential accumulation of GUS in the BS, but that substitution of the guanine at position 3 and thymine at position 6 abolished BS accumulation of GUS (Figure 2B). Similarly, three and five base pair substitutions in BSM1b resulted in a decrease of cell specificity (Figure 2B). Based on these results we propose that within the coding region of *NAD-ME1*, two separate sequences separated by a spacer are necessary and sufficient to generate strong expression in BS cells.

**Figure 2:**
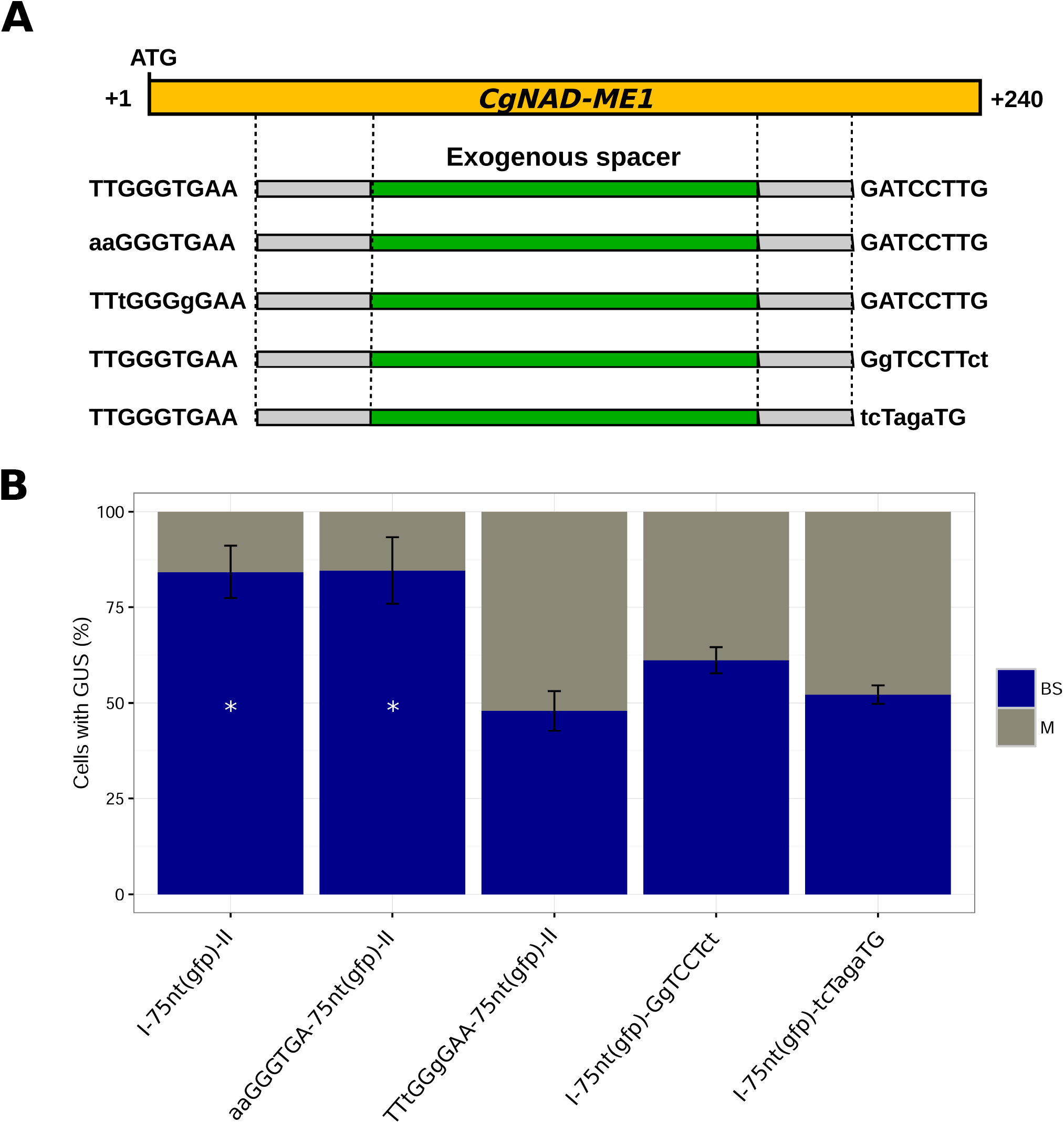
**Two *cis*-elements that are sufficient for preferential accumulation of GUS in the bundle sheath.** Non-mutated and mutated versions of BSM1a and BSM1b flanked by 75 nucleotides derived from GFP were translationally fused to *uidA* encoding GUS and placed under control of the CaMV35S promoter **(A).** The percentage of cells containing GUS after microprojectile bombardment of *G. gynandra* leaves **(B).** Error bars denote the standard error. * represents statistically significant differences with P-values <0.05 and CI = 95% determined by a one-tailed t test.

### BSM1a and 1b specify the spatial patterning of additional genes

Although thousands of genes are differentially expressed between M and BS cells of C_4_ plants, to our knowledge no DNA motifs that determine the patterning of more than one gene in BS cells have been identified. To test whether BSM1a and BSM1b operate more widely to generate preferential expression in BS cells, the coding sequences of other genes were scanned using FIMO (Grant et al., 2011). Sequences similar to BSM1a and BSM1b in genes annotated mitochondrial *MALATE DEHYDROGENASE (mMDH)* and *GLYCOLATE OXIDASE 1 (GOX1)* were identified. In both cases, fragments from *mMDH* and *GOX* containing the two motifs were sufficient to drive BS accumulation of GUS in *G. gynandra* (Figure 3A), and when they were deleted preferential accumulation in BS cells was lost (Figure 3A). The identification of BSM1a and BSM1b in these additional genes allowed consensus sequences to be defined (Figure 3B). These data imply that the DNA sequences defined by BSM1a and BSM1b form the basis of a regulon that operates through conserved *cis*-elements located in the exons of multiple genes to generate preferential expression in BS cells of C_4_ leaves. Altogether these results suggest multiple gene families involved in C_4_ photosynthesis and photorespiration have been recruited into the BS (Figure 3C) using a regulatory network based on these two motifs.

**Figure 3:**
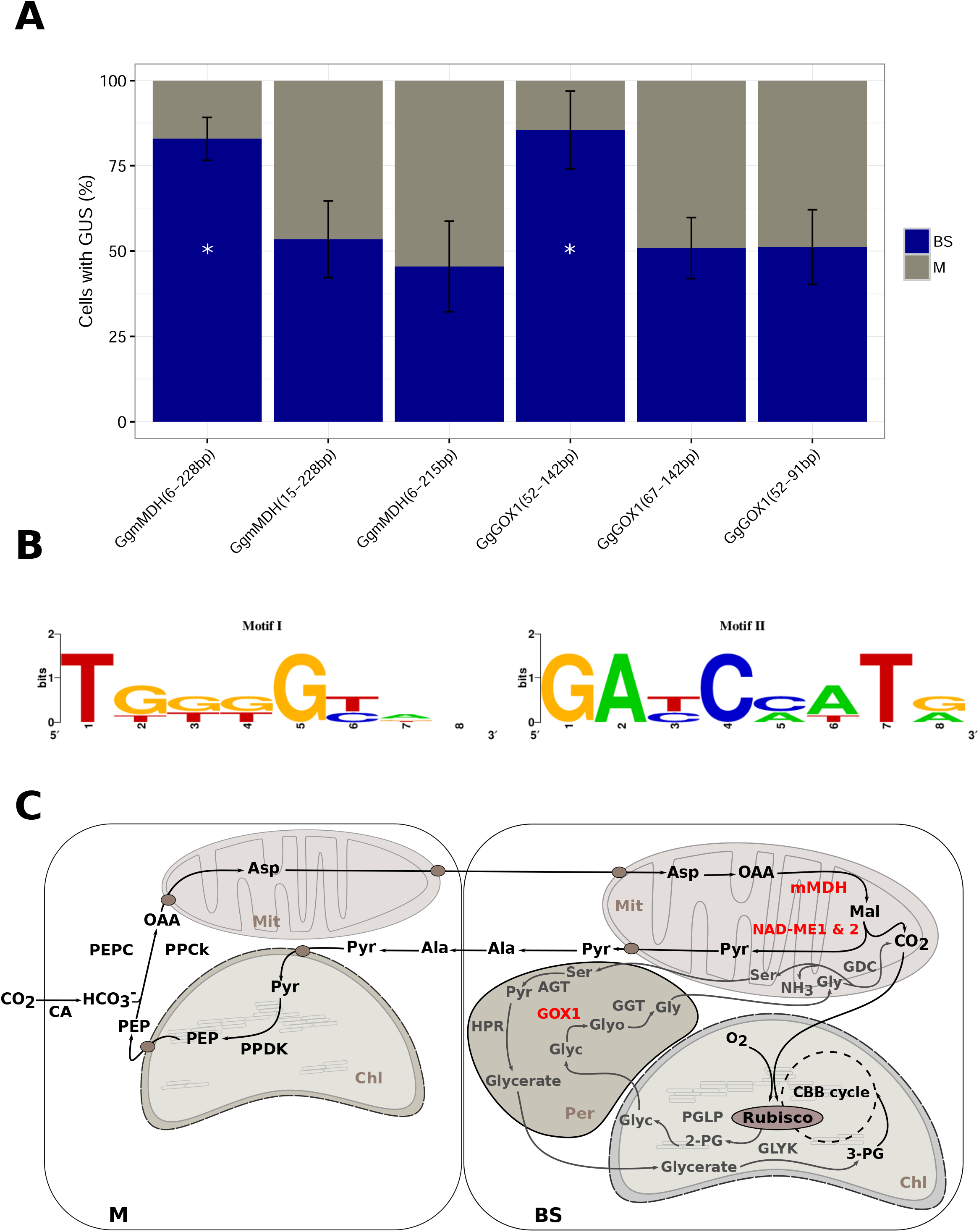
**BSM1a and BSM1b drive the expression of additional genes in C**_**4**_ **and photorespiration pathways.** Sequences similar to BSM1a and BSM1b were predicted to be present in coding sequences of *mMDH* and *GOX1* genes of *G. gynandra.* Deleting the motifs resulted in the loss of preferential accumulation of GUS in the BS **(A).** A consensus sequence for both motifs was defined based on *NAD-ME1, mMDH* and *GOX1* versions of the motifs **(B).** BSM1a and BSM1b coordinate BS gene expression of multiple gene families (highlighted in red) relevant to C_4_ photosynthesis and photorespiration **(C).** Error bars denote the standard error. * represents statistically significant differences with P-values <0.05 and CI = 95% determined by a one-tailed t test.

### BSM1a and BSM1b are ancient and conserved within land plants

Using the sequences that define BSM1a and BSM1b (Figure 3B), and the minimum and maximum distance that can separate them, *NAD-ME* genes from *G. gynandra* and the closely related C_3_ species *A. thaliana* were assessed. Similar sequences close to the predicted translational start site of *GgNAD-ME2, AtNAD-ME1* and *AtNAD-ME2* were identified (Figure 4A). BSM1a is located in the predicted mitochondrial transit peptide and its position varies relative to the translational start site. It is noteworthy that compared with *AtNAD-ME2* and *CgNAD-ME1 & 2*, BSM1a is found on the opposite DNA strand in *AtNAD-ME1*, further supporting the notion that BS preferential expression is mediated by a transcription-based mechanism. BSM1b is located in the mature processed protein and its position appears invariant (Figure 4A). When either of these motifs was removed from *GgNAD-ME2, AtNAD-ME* or *AtNAD-ME2* preferential accumulation in BS cells was lost (Figure 4B). These data indicate that the consensus sequences defined by BSM1a and BSM1b from these eight genes (Figure 4C) are necessary and sufficient to generate BS expression in the C_4_ leaf.

**Figure 4:**
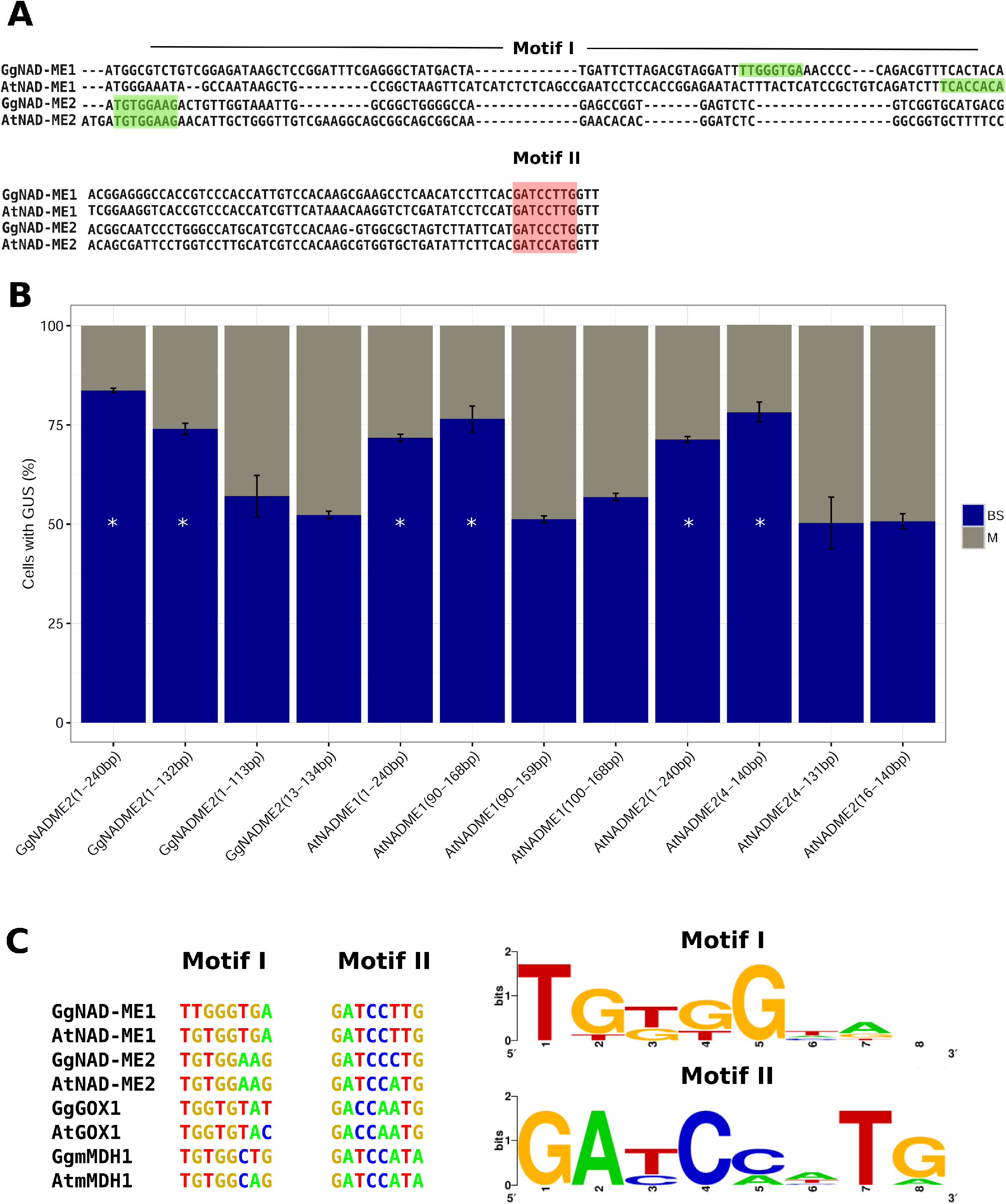
**Functional versions of BSM1a and BSM1b are present in additional *NAD-MEs.*** BSM1a and BSM1b are found in *GgNAD-ME2* and in orthologs of *GgNAD-ME1&2* from the C_3_ species *A. thaliana* **(A).** Translational fusions carrying these fragments confer BS preferential expression in *G. gynandra* leaves. When BSM1a or BSM1b were removed this pattern of GUS was lost **(B).** A consensus sequence generated from all versions of BSM1a and BSM1b tested experimentally **(C).** Error bars denote the standard error. * represents statistically significant differences with P-values <0.05 and CI = 95% determined by a one-tailed t test.

As BSM1a and BSM1b are present in *NAD-ME* genes of C_3_ *A. thaliana*, we next investigated the extent to which these sequences are conserved across 1135 wild inbred *A. thaliana* accessions with genome sequence available. Single nucleotide polymorphism (SNP) data were retrieved (1001 Genomes Consortium, 2016), and analysis showed an unexpectedly high level of conservation with no SNPs detected within either BSM1a or BSM1b (Figure 5A&B). This high level of conservation is consistent with both Motifs acting as “duons” in C_3_ *A. thaliana* as well as C_4_ *G. gynandra*. To investigate whether these motifs are also found more widely in *NAD-ME* genes across the land plant phylogeny, *NAD-ME* gene sequences were retrieved from 44 species in Phytozome (v10.1, www.phytozome.org) and analysed for the presence of BSM1a and BSM1b (Figure S4). All dicotyledons contained at least one *NAD-ME* gene carrying the sequences that define BSM1a and BSM1b (Figure 5C). In the monocotyledons, BSM1a was completely conserved in rice, *Brachypodium* and *Panicum.* Although BSM1b showed one nucleotide substitution in all monocotyledenous genomes available it appears more ancient as it is conserved in spikemoss and moss (Figure 5C, S4). The hypothesis that BSM1b is more ancient is supported by the finding that a version with one nucleotide substitution was also found in the chlorophyte algae *C. reinhardtii* (Figure S4). It was also noticeable that both BSM1a and BSM1b are highly conserved in *GOX1* and *MDH* genes in land plants, and that BSM1b appears more ancient as it is found in all *GOX1* genes from all land plants and even in the chlorophyte algae. It is possible that BSM1a found in land plants is derived from the *MDH* genes of the algae, as it is observed in *C. subellipsoidea* and *M. pusilla*, two members of the chlorophyta. Comparing the sequence of BSM1a and BSM1b in *NAD-ME, MDH* and *GOX1* indicates that BSM1b is less variant, but in both cases, their conservation implies an ancient role across the plant kingdom that likely is derived from the algal ancestor.

**Figure 5:**
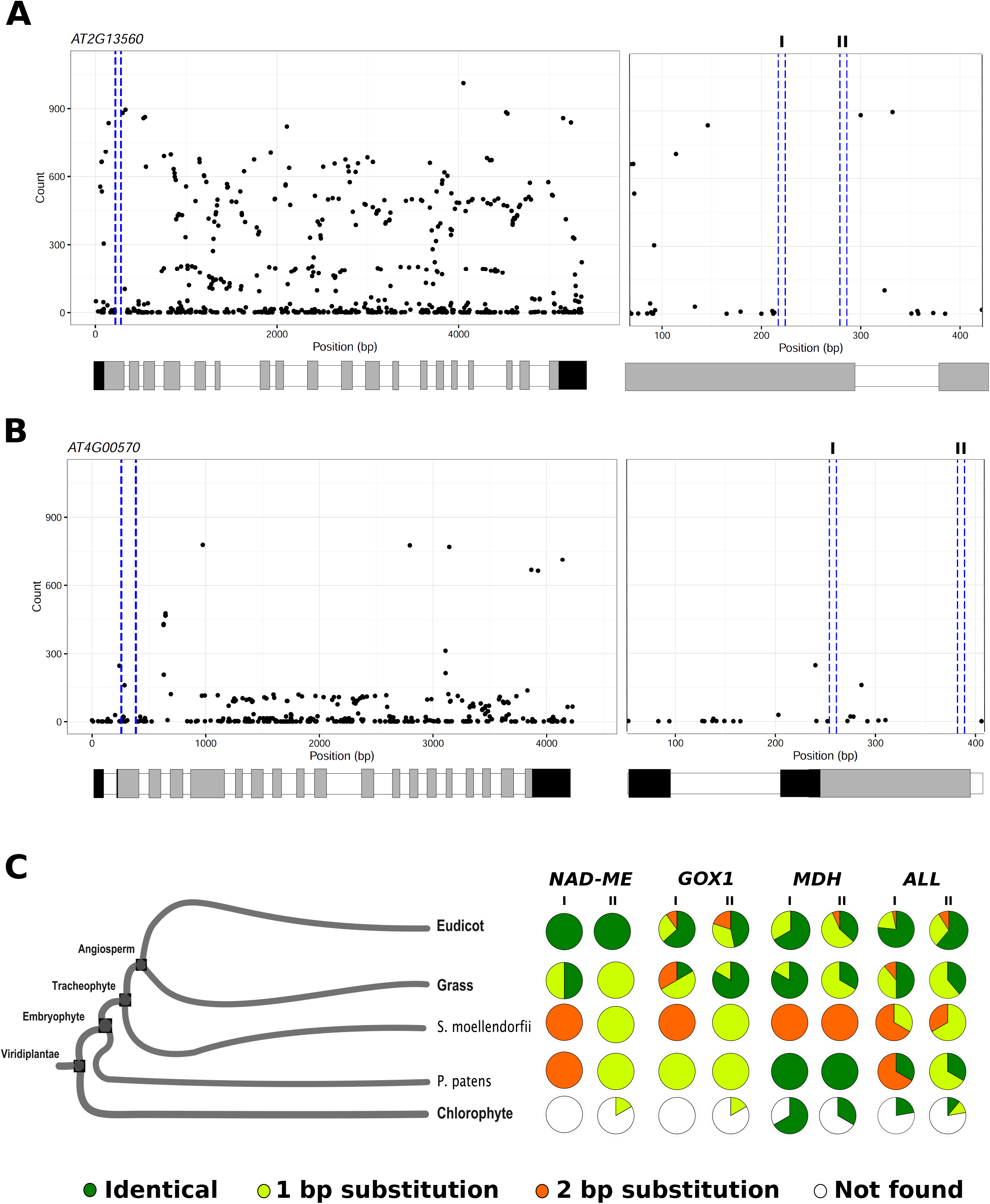
**BSM1a and BSM1b are highly conserved in land plants.** Single nucleotide polymorphisms (SNP) in *AtNAD-ME1* **(A)** and *AtNAD-ME2* **(B)** genes from 1135 wild inbred *A. thaliana* accessions. On the left, the position of BSM1a and BSM1b are highlighted by dashed blue lines, UTRs, exons and introns are denoted by black, grey and white bars respectively on the X-axis. To the right an expanded area representing exon 1, intron 1 and exon 2 is shown, with BSM1a and BSM1b marked within the blue dashed lines. For both genes, no SNP were detected in either motif. The presence of each motifs was investigated in gene sequences of *NAD-ME1, mMDH* and *GOX1* retrieved from 44 species in Phytozome (v10.1). Each Pie-chart shows the percentage of motif instances that were identical (green), or had 1 base pair (yellow), 2 base pair (orange) substitutions or no similarity (white) detected.

### Widespread use of genic DNA for spatial patterning of gene expression

Although genome-wide analysis of transcription factor binding sites indicates a significant amount of binding occurs within genes, to our knowledge, there is little functional knowledge confirming the importance of such sites. Having functionally defined BSM1a and BSM1b as being important for patterning gene expression in the C_4_ leaf, the extent to which other genes important for the C_4_ pathway are regulated by sequences within the gene rather than the promoter was investigated. Based on *de novo* transcriptome assemblies, transcribed regions of twelve genes recruited into C_4_ photosynthesis in *G. gynandra* were cloned, with 3’ and 5’ ends verified using 3’RACE and genome walking (Supplementary Files 1 and 2). The gene sequences were placed under control of the constitutive CaMV35S promoter, and fused to *uidA* encoding the β-glucoronidase (GUS) reporter. Combined with publically available datasets for *PPDK* and *CA* (Kajala et al 2011, Williams et al., 2016) this approach showed that preferential expression of genes encoding C_4_ proteins in either M or BS cells is commonly driven by elements within their coding sequences (Figure 6). *CA2, CA4, PPCk1* and *PPDK* gene sequence without their endogenous promoters all led to preferential accumulation of GUS in M cells, whereas *MDH1, NAD-ME1* and *NAD-ME2* caused preferential accumulation of GUS in the BS. *Rubisco Activase (RCA)* coding sequence drove a small but significant increase in the number of BS accumulating GUS (Figure 6A) (p-value <0.05, CI 95%). These data indicate that regulatory elements within genic sequence impact on cell preferential expression in the majority of genes recruited into the core C_4_ pathway.

**Figure 6:**
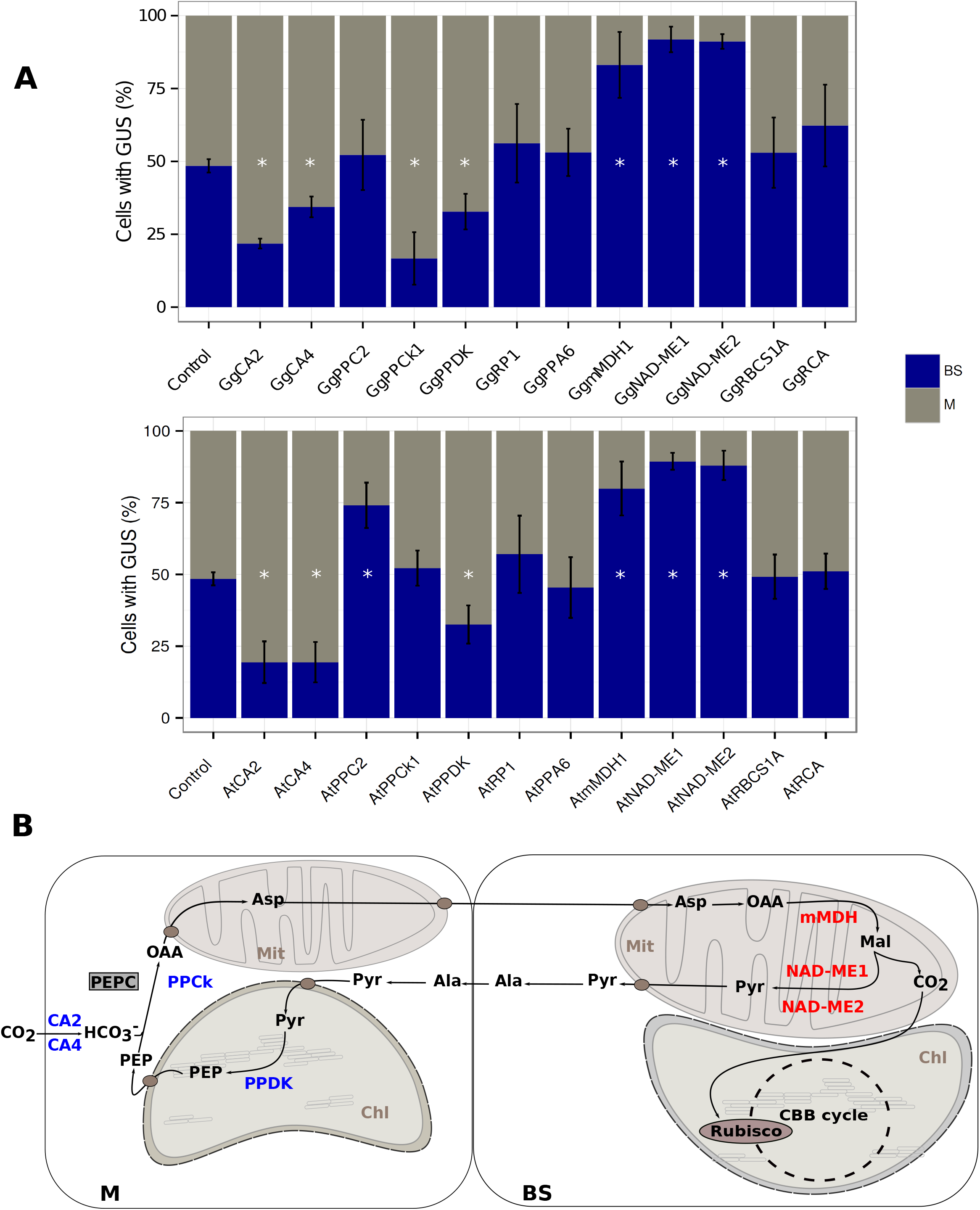
**Pre-existing intragenic regulatory sequences play a major role controlling C**_**4**_ **photosynthesis genes.** Coding sequences encoding for core proteins of the C_4_ pathway from *G. gynandra* together with orthologs from *A. thaliana* were translationally fused to *uidA* and placed under control of the CaMV35S promoter. After introduction into *G. gynandra* leaves by microprojectile bombardment mesophyll preferential expression of *CA2, CA4, PPDK* and *PPCk*, together with Bundle Sheath preferential expression of *mMDH, NAD-ME1* and *NAD-ME2* were observed **(A).** With the exception of *PPCk* these regulatory elements are conserved in orthogues from *A. thaliana* **(B).** The contribution of intragenic sequences controlling gene regulation of the C_4_ pathway is summarized in **(C),** *CA2, CA4, PPDK* and *PPCk* (blue) and *mMDH, NAD-ME1&2* (red) denote genes where intragenic sequences control cell preferential gene expression. Error bars denote the standard error. * represents statistically significant differences with P-values <0.05 and CI = 95% determined by a one-tailed t test.

In some cases, cell-preferential expression of C_4_ genes has evolved from regulatory elements found in ancestral C_3_ species (Kajala, et al., 2011; Brown et al., 2011, Williams et al., 2013). To investigate the extent to which genic sequence from ancestral C_3_ species contain regulatory elements sufficient for expression in either M or BS cells, orthologues to each of the C_4_ genes were cloned from *A. thaliana*, placed under the same reporter system and tested by microprojectile bombardment. This showed that with the exception of *AtPPCk1*, the orthologous genes from C_3_ *A. thaliana* contained regulatory elements in coding sequence that can specify spatial patterning of gene expression in the C_4_ leaf of *G. gynandra* (Figure 6). Overall, these data indicate that spatial patterning of gene expression in the C_4_ leaf is largely derived from regulatory elements present in coding sequences of genes found in the ancestral C_3_ state. In the case of those defined at the nucleotide level in *NAD-ME1, NAD-ME2, MDH* and *GOX1*, these elements appear highly conserved and therefore ancient within land plants.

## Discussion

The data presented here, combined with previous reports (Brown et al. 2011; Kajala et al. 2011; Williams et al. 2016) portray an overview of the contribution that untranslated regions (UTRs) and coding sequences make to the generation of cell-specific gene expression in leaves of C_4_ *G. gynandra.* Eight of the eleven core C_4_ cycle genes possess regulatory elements in their transcript sequences that are sufficient for preferential accumulation in either M or BS cells of the C_4_ leaf. These data strongly imply that, in addition to promoters being involved in generating cell-specificity in C_4_ leaves (Gowik et al., 2004; Sheen, 1999), coding sequences and UTRs play a widespread role in the preferential accumulation of C_4_ transcripts to either M or BS cells. It remains to be seen whether this high degree of regulation from genic sequence is a common phenomenon in C_4_ leaves of species other than *G. gynandropsis*, or whether it is critical for the spatial control of gene expression in other tissues and other species. However, as genome-wide studies of transcription factor recognition sites in organisms as diverse as *A. thaliana* and human cells (Stergachis et al., 2013; Sullivan et al., 2014) have reported significant binding occurs in genic sequence, we anticipate many more examples of spatial regulation of gene expression being associated with *cis-*elements outside of promoter sequences.

The accumulation of *GgNADME1, GgNADME2, mMDH* and *GOX1* transcripts in BS cells is dependent on the co-operative function of two *cis*-elements that are separated by a spacer sequence, all of which are located in the first exon of these genes. These motifs are conserved both in their sequences, but also in the number of nucleotides that separates them. In orthologous *NAD-ME* genes from C_3_ *A. thaliana*, which diverged from the Cleomaceae ~38 million years ago (Schranz and Mitchell-Olds, 2006; Couvreur et al., 2010), although these motifs are present, they are not sufficient to generate cell preferential expression in the C_3_ leaf (Brown et al. 2011). This finding indicates that for *NAD-ME* genes to be preferentially expressed in BS cells of C_4_ plants, a change in the behaviour of one or more *trans*-factors was a fundamental event. At least in *G. gynandropsis*, evolution appears to have repeatedly made use of *cis*-elements that exist in genes of C_3_ species that are orthologous to those recruited into C_4_ photosynthesis (Brown et al., 2011; Kajala et al., 2012; Williams, Burgess et al., 2016). The alteration in *trans*-factors such that they recognise ancestral elements in *cis* in the M or BS therefore appears to be an important and common mechanism associated with evolution of the highly complex C_4_ system.

BSM1a and BSM1b, which we defined first in the *GgNAD-ME1* gene, are also present and operational in the *GgNAD-ME2, GgMDH1* and *GgGOX1* genes. If, as seems likely, these motifs are recognised by the same *trans*-factors to generate preferential expression of all of these genes in the C_4_ BS, this finding also identifies a mini-regulon that during evolution could have recruited at least four genes simultaneously into specialised roles in the BS. By combining Flux Balance Analysis constrained by a model of carbon fixation, it has previously been proposed that upregulation and preferential expression of multiple genes of the C_4_ cycle would be required to balance nitrogen metabolism between M and BS cells (Mallmann et al., 2014). The presence of BSM1a and BSM1b in at least four genes from *G. gynandropsis* provides a mechanism that may have facilitated this patterning of multiple genes during the evolution of C_4_ photosynthesis.

The dual role of exons in protein coding as well as the regulation of gene expression has received significant attention in vertebrates (Lang et al., 2005; Nguyen et al., 2007; Goren et al., 2006; Tumpel et al., 2008; Dong et al., 2010, Stergachis et al., 2013). Although, 11% of transcription factor binding sites are located in exonic sequence in *A. thaliana* (Sullivan et al., 2014), to our knowledge, the identification of BSM1a and BSM1b represents the first functional evidence for *cis*-elements in plant exons. The fact that these motifs are present in C_3_ *A. thaliana*, and in fact, also found in the genomes of many land plants and some chlorophyte algae, indicates that these duons play ancient and conserved roles in photosynthetic organisms. The role of such regulatory elements within coding sequences has previously been proposed to be associated with constraints on both protein coding function and codon bias. For example, mutation to these *cis-*elements could be deleterious to both the correct function of the protein, but also to codon usage and so translational efficiency (Robinson et al., 1984; Tuller et al., 2010; Nakahigashi et al., 2014). If this is the case, BSM1a and BSM1b could be highly conserved across deep phylogeny because of strong positive selection pressure on these elements due to impact on translation, and this conservation is then co-opted to also regulate transcription during the evolution of C_4_ photosynthesis to generate cell-specific gene expression. Establishing the role of BSM1a and BSM1b in C_3_ plants would provide insight into the extent to which their role has altered during the transition from C_3_ to C_4_ photosynthesis.

Duons under strong selection pressure may represent a rich resource of *cis*-elements upon which the C_4_ pathway has evolved. Although C_4_ photosynthesis is a complex trait that requires multiple changes to gene expression, the repeated recurrence of C_4_ species across multiple plant lineages suggests that a relatively low number of changes may be required to acquire the C_4_ syndrome (Sinha & Kellogg, 1996; Hibberd et al., 2008; Westhoff & Gowik, 2010). A single C_4_ master switch has been proposed (Westhoff & Gowik, 2010) but despite multiple comparative transcriptomic studies (Brautigam et al., 2011; Aubry et al., 2014; Kulahoglu et al., 2014), there is as yet no evidence for it. Given the repeated and highly convergent evolution of the C_4_ pathway, as well as evidence that separate lineages can arrive at the C_4_ state via different routes (Williams et al., 2013), it appears more plausible that C_4_ photosynthesis made use of a number of gene sub-networks. This is now supported by a number of findings. First, just as core photosynthesis genes encoding the light harvesting complexes and Calvin-Benson-Bassham cycle are regulated by light, the vast majority of genes that encode proteins of the C_4_ cycle in C_3_ *A. thaliana* are also regulated by light signalling, yet, during the evolution of C_4_ photosynthesis there was a significant gain of responsiveness to chloroplast signalling (Burgess et al., 2016). Second, it has been suggested that evolution of the C_4_ pathway is associated with the recruitment of developmental motifs into leaves that in C_3_ species operate in roots (Kulahoglu et al., 2014). Lastly, the identification of the *cis*-element MEM2 (Williams, Burgess et al., 2016), which controls preferential expression of multiple genes in C_4_ M cells, and now BSM1a and BSM1b in four different genes that are strongly expressed in BS cells, indicates that that C_4_ evolution has made use of small-scale recruitment of gene sub-networks in both cell-types.

## Methods

### Growth of plant material and production of reporter constructs

Sterile *G. gynandra* seed was sown directly from intact pods and germinated on moist filter papers in the dark at 30°C for 24 h. Seedlings were then transferred to Murashige and Skoog (MS) medium with 1% (w/v) sucrose and 0.8% (w/v) agar (pH 5.8) and grown for a further 13 days in a growth room at 22°C and 200 µmol m^−2^ s^-1^ photon flux density (PFD) with a photoperiod of 16 h light.

*G. gynandropsis* mRNA sequences were predicted from a *de novo* assembled transcriptome and UTRs were verified by 3’ RACE (Supplementary File I) and genome walking (Supplementary File 2). *A. thaliana* cDNA sequences were extracted from Phytozome v10.1. Reporter constructs were generated by ligation of the fragment of interest with a modified reporter cassette containing the Cauliflower Mosaic Virus 35S promoter (pCaMV35S), 13 bp of its 5’UTR, the *uidA* gene (encoding GUS), and the *nosT* terminator sequence (Brown et al., 2011). Vectors were assembled in this cassette using Gibson assembly (Gibson et al., 2009) (Supplementary Table I). Site-directed mutagenesis was performed using the Quickchange method.

### Microprojectile bombardment and production of stable transformants

350 ng M-17 tungsten particles (1.1-µm diameter; Bio-Rad) were washed with 100% (v/v) ethanol and resuspended in ultrapure water. 1.5 µg of plasmid DNA was mixed with the tungsten particles while vortexing at slow speed. After addition of the DNA, 50 µL 2.5 M calcium chloride (Fisher Scientific) and 10 µL 100 mM spermidine (Sigma-Aldrich) were added to the particle suspension to facilitate binding of DNA to the particles. The tungsten-DNA suspension was incubated for 10 min on ice, with frequent agitation to prevent pelleting. Particles were then washed and resuspended in 100 µL 100% (v/v) ethanol. 10 µL aliquots of tungsten-DNA were transferred to plastic macrocarriers (Bio-Rad) and allowed to dry for 3 minutes at room temperature. Three macrocarriers were used for each transformation. Following bombardment with a Bio-Rad PDS-1000/He particle delivery system, seedlings were placed upright in a sealed Petri dish, with the base of their stems immersed in 0.5x MS medium and incubated in a growth room at 22°C and 150 µmol m^−2^ s^-1^ PFD with a photoperiod of 16 h light for 48 h, prior to GUS staining. Stable plant transformation was performed by introducing constructs into *G.gynandra* via *Agrobacterium tumefaciens* LBA4404 as described previously (Newell et al., 2009). Plant tissue, after bombardment or stable transformation, was GUS stained (0.1 M Na2HPO4 pH7.0, 0.5 mM K ferricyanide, 0.5 mM K ferrocyanide, 0.06% v/v Triton X-100, 10 mM Na_2_EDTA pH8.0, 1mM X-gluc) at 37°C for 6-16 h and then fixed in a 3:1 solution of ethanol to acetic acid at room temperature for 30 min. Chlorophyll was cleared with 70% (v/v) ethanol and tissue treated with 5% (w/v) NaOH at 37°C for 2 h. M and BS cells containing GUS were identified and counted using phase-contrast microscopy. At least 50 cells were counted per construct in each experiment, and for each construct, three independent experiments were conducted (Supplementary Table II).

### *cis*-Element prediction and localization

*De novo* motif prediction was performed using the Multiple Em for Motif Elucidation (MEME) suite v.4.8.1 with the following parameters: *meme sequences.fa -dna -oc. -nostatus -time 18000 -maxsize 60000 -mod oops -nmotifs 3 -minw 7 -maxw 9 -revcomp.* To scan for motif instances across various datasets FIMO was used with the following parameters: *fimo --oc. --verbosity 1 --thresh 0.1 motifs, meme sequences.fa.* Only hits located within the first 550 bp, allowing a spacing between the motifs of 35 to 550 bp were accepted.

## Figure legends

**Figure S1: Transformation of *G. gynandra* M and BS cells by microprojectile bombardment.** Leaves of *G. gynandra* arranged concentrically prior to bombardment **(A).** Representative GUS stained *G. gynandra* leaf transformed with *pCaMV35s:GgNAD-ME1(25-240bp)::gfp/uidA::nosT* **(B).** Mesophyll cells **(C)** and Bundle Sheath cells (D, black arrows) stained with GUS after bombardment. Scale bars represent 100 µm.

**Figure S2: Pixel intensity of stable transgenic lines.** Pixel intensities across regions of the leaf containing mesophyll and bundle sheath. Data are derived from GUS stained leaves from three independent transgenic lines. Data are presented as histograms for whole datasets and dots that represent single measurements. At least 20 measurements were made per transgenic line.

**Figure S3: Topological requirements for BSM1a and BSM1b function.** Summary of the constructs used in this experiment. BSM1a and BSM1b were separated by 0, 21, 240, 347, 413, 550 and 999 base pairs derived from the gene encoding Green Fluorescent Protein (GFP) **(A).** Percentage of cells containing GUS after microprojectile bombardment of *G. gynandra* leaves. Bars represent the percentage of stained cells in Bundle Sheath (BS - blue) and mesophyll (M-grey) cells. Error bars denote the standard error of the mean. * represents statistically significant differences with P-values <0.05 and CI = 95% determined by a one-tailed t test.

**Figure S4: Location of BSM1a and BSM1b across the land plant phylogeny.** BSM1a and BSM1b in *NAD-ME, mMDH* and *GOX1* coding sequences retrieved from 44 species in Phytozome (v10.1). Green dots represent identical versions of the motifs while yellow and orange dots denote alternative versions with one or two substitutions respectively.

**Supplementary File 1:** FASTA sequences from Rapid Amplification of cDNA ends used to verify 3’ UTR sequences of C_4_ genes from *G. gynandra.*

**Supplementary File 2:** FASTA sequences from Genome Walking experiments used to verify 5’ ends of C_4_ gene sequences from *G. gynandra.*

**Supplementary Table I:** Primer sequences used in generation of constructs, 3’ RACE experiments and Genome Walking.

**Supplementary Table II:** Total cell counts for the microprojectile bombardment experiments.

## References

Akyildiz, M., Gowik, U., Engelmann, S., Koczor, M., Streubel, M., & Westhoff, P. (2007). Evolution and Function of a cis-Regulatory Module for Mesophyll-Specific Gene Expression in the C 4 Dicot Flaveria trinervia. The Plant, 19(November), 3391–3402. http://doi.org/10.1105/tpc.107.053322

Anbar, A. D., Duan, Y., Lyons, T. W., Arnold, G. L., Kendall, B., Creaser, R. A., … Hayes, M. (2007). A Whiff of Oxygen before the Great Oxidation Event? Source: Science, New Series Nature Science Philos. Trans. R. Soc London Ser. B Science Rapid Commun. Mass Spectrom S. Ono et Ai Earth Planet Sei Lett. N. J. Beukes, Sediment Geol Econ. Geol, 317(84), 1903–1906. http://doi.org/10.1126/science.1140325

Aubry, S., Kelly, S., Kümpers, B., & Smith-Unna, R. (2014). Deep evolutionary comparison of gene expression identifies parallel recruitment of trans-factors in two independent origins of C 4 photosynthesis. PLoS Genetics. Retrieved from http://jurnals.plos.org/plosgenetics/article?id=10.1371/journal.pgen.1004365

Bailey, T. L., Boden, M., Buske, F. A., Frith, M., Grant, C. E., Clementi, L., … Noble, W. S. (2009). MEME S UITE: tools for motif discovery and searching. Conflict, 37(May), 202–208. http://doi.org/10.1093/nar/gkp335

Bauwe, H., Hagemann, M., & Fernie, A. R. (2010). Photorespiration: players, partners and origin. Trends in Plant Science, 15(6), 330–6. http://doi.org/10.1016/j.tplants.2010.03.006

Bräutigam, A., Kajala, K., Wullenweber, J., & Sommer, M. (2011). mRNA Blueprint for C Photosynthesis Derived from Comparative Transcriptomics of Closely Related C### and C### Species. Plant Physiology. Retrieved from http://agris.fao.org/agris-search/search.do?recordID=US201301934603

Brown, N. J., Newell, C. a, Stanley, S., Chen, J. E., Perrin, A. J., Kajala, K., & Hibberd, J. M. (2011). Independent and parallel recruitment of preexisting mechanisms underlying C photosynthesis. xScience (New York, N.Y.), 331(6023), 1436–1439. http://doi.org/10.1126/science.1201248

Burgess, S. J., Granero-moya, I., Grangé-guermente, M. J., Boursnell, C., Terry, M. J., & Hibberd, J. M. (2016). Ancestral light and chloroplast regulation form the foundations for C 4 gene expression. Nature Plants. http://doi.org/10.1038/nplants.2016.161

Consortium, 1001 Genomes. (2016). 1,135 Genomes Reveal the Global Pattern of Polymorphism in Arabidopsis thaliana. Cell. Retrieved from http://www.sciencedirect.com/science/article/pii/S0092867416306675

Couvreur, T. L. P., Franzke, A., Al-Shehbaz, I. A., Bakker, F. T., Koch, M. A., & Mummenhoff, K. (2010). Molecular phylogenetics, temporal diversification, and principles of evolution in the mustard family (Brassicaceae). Molecular Biology and Evolution, 27(1), 55–71. http://doi.org/10.1093/molbev/msp202

Crooks, G., Hon, G., Chandonia, J., & Brenner, S. (2004). NCBI GenBank FTP Site\nWebLogo: a sequence logo generator. Genome Res, 14, 1188–1190. http://doi.org/10.1101/gr.849004.1

Dong, X., Navratilova, P., Fredman, D., Drivenes, Becker, T. S., & Lenhard, B. (2009). Exonic remnants of whole-genome duplication reveal cis-regulatory function of coding exons. Nucleic Acids Research, 38(4), 1071–1085. http://doi.org/10.1093/nar/gkp1124

Edwards, G. E., Franceschi, V. R., & Voznesenskaya, E. V. (2004). SINGLE-CELL C4 PHOTOSYNTHESIS VERSUS THE DUAL-CELL (KRANZ) PARADIGM. Annual Review of Plant Biology, 55(1), 173–196. http://doi.org/10.1146/annurev.arplant.55.031903.141725

Fankhauser, N., Aubry, S. (2016). Post-transcriptional regulation of photosynthetic genes is a key driver of C4 leaf ontogeny. Journal of Experimental Botany, erw386. http://jxb.oxfordiournals.org/content/early/2016/10/17/jxb.erw386.abstract

Fontaine, V., Hartwell, J., Jenkins, G., & Nimmo, H. (2002). Arabidopsis thaliana contains two phosphoenolpyruvate carboxylase kinase genes with different expression patterns. Plant, Cell & Environment, 25, 115–122. http://doi.org/10.1046/j.0016-8025.2001.00805.x

Furbank, R. T. (2011). Evolution of the C4 photosynthetic mechanism: Are there really three C4 acid decarboxylation types? Journal of Experimental Botany, 62(9), 3103–3108. http://doi.org/10.1093/jxb/err080

G, L.-M. (2006). Comparative analysis identifies exonic splicing regulatory sequences-the complex definition of enhancers and silencers. Mol. Cell, 22, 769. Retrieved from http://www.sciencedirect.com/science/article/pii/S1097276506003005

Gibson, D. G., Young, L., Chuang, R.-Y., Venter, J. C., Hutchison, C. a, Smith, H. O., … America, N. (2009). Enzymatic assembly of DNA molecules up to several hundred kilobases. Nature Methods, 6(5), 343–5. http://doi.org/10.1038/nmeth.1318

Gowik, U., Burscheidt, J., Akyildiz, M., Schlue, U., Koczor, M., Streubel, M., & Westhoff, P. (2004). cis-Regulatory Elements for Mesophyll-Specific Gene Expression in the C 4 Plant Flaveria trinervia, the Promoter of the C 4 Phosphoenolpyruvate Carboxylase Gene. The Plant Cell, 16(May), 1077–1090. http://doi.org/10.1105/tpc.019729.C

Grant, C. E., Bailey, T. L., & Noble, W. S. (2011). FIMO: Scanning for occurrences of a given motif. Bioinformatics, 27(7), 1017–1018. http://doi.org/10.1093/bioinformatics/btr064

Hatch, M. D., & Slack, C. R. (1966). Photosynthesis by sugar-cane leaves. A new carboxylation reaction and the pathway of sugar formation. The Biochemical Journal, 101(1), 103–11. Retrieved from http://www.pubmedcentral.nih.gov/articlerender.fcgi?artid=1270070&tool=pmcentrez&rendertype=abstract

Hibberd, J. M., & Covshoff, S. (2010). The regulation of gene expression required for C4 photosynthesis. Annual Review of Plant Biology, 61(1), 181–207. http://doi.org/doi:10.1146/annurev-arplant-042809-112238

Hibberd, J. M., Sheehy, J. E., & Langdale, J. A. (2008). Using C 4 photosynthesis to increase the yield of rice - rationale and feasibility. Current Opinion in Environmental Sustainability, 11, 4–7. http://doi.org/10.1016/j.pbi.2007.11.002

Kajala, K., Williams, B. P., Brown, N. J., Taylor, L. E., & Hibberd, J. M. (2011). Multiple Arabidopsis genes primed for direct recruitment into C4 photosynthesis. Plant Journal, 69, 47–56. Retrieved from http://onlinelibrary.wiley.com/doi/10.1111/j.1365-313X.2011.04769.x/full

Kausch, a P., Owen, T. P., Zachwieja, S. J., Flynn, a R., & Sheen, J. (2001). Mesophyll-specific, light and metabolic regulation of the C4 PPCZm1 promoter in transgenic maize. Plant Molecular Biology, 45(1), 1–15. http://doi.org/10.1023/A:1006487326533

Kim, D., Pertea, G., Trapnell, C., Pimentel, H., Kelley, R., & Salzberg, S. L. (2013). TopHat2: accurate alignment of transcriptomes in the presence of insertions, deletions and gene fusions. Genome Biology, 14(4), R36. http://doi.org/10.1186/gb-2013-14-4-r36

Krebbers, E., Seurinck, J., Herdies, L., Cashmore, A. R., & Timko, M. P. (1988). Four genes in two diverged subfamilies encode the ribulose-1,5-bisphosphate carboxylase small subunit polypeptides of Arabidopsis thaliana. Plant Molecular Biology, 11(6), 745–759. http://doi.org/10.1007/BF00019515

Külahoglu, C., Denton, A. K., Sommer, M., Maß, J., Schliesky, S., Wrobel, T. J., … Weber, A. P. M. (2014). Comparative transcriptome atlases reveal altered gene expression modules between two Cleomaceae C3 and C4 plant species. The Plant Cell, 26(8), 3243–60. http://doi.org/10.1105/tpc.114.123752

Lang, G., Gombert, W. M., & Gould, H. J. (2005). A transcriptional regulatory element in the coding sequence of the human Bcl-2 gene. Immunology, 114, 25–36. http://doi.org/10.1111/j.1365-2567.2004.02073.x

Li, B., Ruotti, V., Stewart, R. M., Thomson, J. A., & Dewey, C. N. (2009). RNA-Seq gene expression estimation with read mapping uncertainty. Bioinformatics, 26(4), 493–500. http://doi.org/10.1093/bioinformatics/btp692

Mallmann, J., Heckmann, D., Bräutigam, A., Lercher, M. J., Weber, A. P. M., Westhoff, P., & Gowik, U. (2014). The role of photorespiration during the evolution of C^4^ photosynthesis in the genus Flaveria. eLife, 2014(3). http://doi.org/10.7554/eLife.02478

Nakahigashi, K., Takai, Y., Shiwa, Y., Wada, M., Honma, M., Yoshikawa, H., … Mori, H. (2014). Effect of codon adaptation on codon-level and gene-level translation efficiency in vivo. BMC Genomics, 15(1), 1115. http://doi.org/10.1186/1471-2164-15-1115

Newell, C. A., Brown, N. J., Liu, Z., Pflug, A., Gowik, U., Westhoff, P., & Hibberd, M. (2010). Agrobacterium tumefaciens-mediated transformation of Cleome gynandra L., a C 4 dicotyledon that is closely related to Arabidopsis thaliana. Journal of Experimental Botany, 61(5), 1311–1319. http://doi.org/10.1093/jxb/erq009

Nguyen, M. Q., Zhou, Z., Marks, C. A., Ryba, N. J. P., & Belluscio, L. (2007). Prominent Roles for Odorant Receptor Coding Sequences in Allelic Exclusion. Cell, 131(5), 1009–1017. http://doi.org/10.1016/j.cell.2007.10.050

Nomura, M., Sentoku, N., Nishimura, A., Lin, J. H., Honda, C., Taniguchi, M., … Matsuoka, M. (2000). The evolution of C4 plants: Acquisition of cis-regulatory sequences in the promoter of C4-type pyruvate, orthophosphate dikinase gene. Plant Journal, 22(3), 211–221. http://doi.org/10.1046/j.1365-313X.2000.00726.x

Ray, D. K., Ramankutty, N., Mueller, N. D., West, P. C., & Foley, J. a. (2012). Recent patterns of crop yield growth and stagnation. Nature Communications, 3, 1293. http://doi.org/10.1038/ncomms2296

Robinson, M., Lilley, R., Little, S., & Emtage, J. (1984). Codon usage can affect efficiency of translation of genes in Escherichia coli. Nucleic Acids. Retrieved from http://nar.oxfordjournals.org/content/12/17/6663.short

Sage, R. F. (2004). Tansley review: The evolution of C4 photosynthesis. New Phytol, 161, 30. Retrieved from http://onlinelibrary.wiley.com/doi/10.1111/j.1469-8137.2004.00974.x/full

Sage, R. F., Wedin, D. a, & Li, M. (1999). The Biogeography of C4 Photosynthesis: Patterns and Controlling Factors. C4 Plant Biology, 313–373. http://doi.org/10.1016/B978-012614440-6/50011-2

Sanchez, R., & Cejudo, F. (2003). Identification and expression analysis of a gene encoding a bacterial-type phosphoenolpyruvate carboxylase from Arabidopsis and rice. Plant Phys/ology, 132(June), 949–957. http://doi.org/10.1104/pp.102.019653.1997

Schranz, M. E., & Mitchell-Olds, T. (2006). Independent ancient polyploidy events in the sister families Brassicaceae and Cleomaceae. The Plant Cell, 18(5), 1152–1165. http://doi.org/10.1105/tpc.106.041111

Schulze, S., Mant, A., Kossmann, J., & Lloyd, J. R. (2004). Identification of an Arabidopsis inorganic pyrophosphatase capable of being imported into chloroplasts. FEBS Letters, 565(1–3), 101–105. http://doi.org/10.1016/j.febslet.2004.03.080

Sheen, J. (1991). Molecular mechanisms underlying the differential expression of maize pyruvate, orthophosphate dikinase genes. The Plant Cell, 3(3), 225–245. http://doi.org/10.1105/tpc.3.3.225

Sheen, J. (1999). C 4 Gene Expression. Annual Review of Plant Physiology and Plant Molecular Biology, 50(1), 187–217. Retrieved from http://www.annualreviews.org/doi/pdf/10.1146/annurev.arplant.50.L187

Sinha, N. R., & Kellogg, E. A. (1996). Parallelism and diversity in multiple origins of C4 photosynthesis in the grass family. American Journal of Botany, 83(11), 1458–1470. http://doi.org/10.2307/2446101

Stergachis, A. B., Haugen, E., Shafer, A., Fu, W., Vernot, B., Reynolds, A., … Stamatoyannopoulos, J. a. (2013). Exonic transcription factor binding directs codon choice and affects protein evolution. Science (New York, N.Y.), 342(6164), 1367–72. http://doi.org/10.1126/science.1243490

Sullivan, A. M., Arsovski, A. A., Lempe, J., Bubb, K. L., Weirauch, M. T., Sabo, P. J., … Stamatoyannopoulos, J. A. (2014). Mapping and dynamics of regulatory DNA and transcription factor networks in A. thaliana. Cell Reports, 8(6), 2015–2030. http://doi.org/10.1016/j.celrep.2014.08.019

Tuller, T., Waldman, Y. Y., Kupiec, M., & Ruppin, E. (2010). Translation efficiency is determined by both codon bias and folding energy. Proceedings of the National Academy of Sciences of the United States of America, 107(8), 3645–50. http://doi.org/10.1073/pnas.0909910107

Tümpel, S., Cambronero, F., Sims, C., Krumlauf, R., & Wiedemann, L. M. (2008). A regulatory module embedded in the coding region of Hoxa2 controls expression in rhombomere 2. Proceedings of the National Academy of Sciences of the United States of America, 105(51), 20077–20082. http://doi.org/10.1073/pnas.0806360105

Viret, J., Mabrouk, Y., & Bogorad, L. (1994). Transcriptional photoregulation of cell-type-preferred expression of maize rbcS-m3: 3’and 5’sequences are involved. Proceedings of the. Retrieved from http://www.pnas.org/content/91/18/8577.short

Werneke, J. M., Zielinski, R. E., & Ogren, W. L. (1988). Structure and expression of spinach leaf cDNA encoding ribulosebisphosphate carboxylase/oxygenase activase. Proceedings of the National Academy of Sciences of the United States of America, 85(3), 787–91. Retrieved from http://www.pubmedcentral.nih.gov/articlerender.fcgi?artid=279640&tool=pmcentrez&rendertype=abstract

Westhoff, P., & Gowik, U. (2010). Evolution of C4 photosynthesis--looking for the master switch. Plant Physiology, 154(2), 598–601. http://doi.org/10.1104/pp.110.161729

Williams, B., Burgess, S., & Reyna-Llorens, I. (2016). An untranslated cis-element regulates the accumulation of multiple C4 enzymes in Gynandropsis gynandra mesophyll cells. The Plant. Retrieved from http://www.plantcell.org/content/28/2/454.short

Williams, B. P., Johnston, I. G., Covshoff, S., & Hibberd, J. M. (2013). Phenotypic landscape inference reveals multiple evolutionary paths to C4 photosynthesis. eLife, 2, e00961. http://doi.org/10.7554/eLife.00961

